# xQTLbiolinks: a comprehensive and scalable tool for integrative analysis of molecular QTLs

**DOI:** 10.1101/2023.04.28.538654

**Authors:** Ruofan Ding, Xudong Zou, Yangmei Qin, Hui Chen, Xuelian Ma, Chen Yu, Gao Wang, Lei Li

**Author notes:** These authors contributed equally (Ruofan Ding, Xudong Zou).

## Abstract

Genome-wide association studies (GWAS) have identified thousands of disease-associated non-coding variants, posing urgent needs for functional interpretation. Molecular quantitative trait loci (xQTLs) such as eQTLs serve as an essential intermediate link between these non-coding variants and disease phenotypes and have been widely used to discover disease-risk genes from many population-scale studies. However, mining and analyzing the xQTLs data presents several significant bioinformatics challenges, particularly when it comes to integration with GWAS data. Here, we developed xQTLbiolinks as the first comprehensive and scalable tool for xQTLs data retrieval, quality controls, and pre-processing from 75 human tissues and cell types. In addition, xQTLbiolinks provided a robust colocalization module through integration with GWAS data. The result generated by xQTLbiolinks can be flexibly visualized or stored in standard R objects that can easily be integrated with other R packages and custom pipelines. We applied xQTLbiolinks to cancer GWAS summary statistics as case studies and demonstrated its robust utility and reproducibility. xQTLbiolinks will profoundly accelerate the interpretation of disease-associated variants, thus promoting a better understanding of disease etiologies. xQTLbiolinks is available at https://github.com/lilab-bioinfo/xQTLbiolinks.

## INTRODUCTION

The explosion of GWAS discovery and applications across multiple disciplines yielded many risk loci mainly located in non-coding regions, prompting the great need for functional interpretation of these variants by revealing the underlying mechanisms and susceptibility genes (1). The large-scale molecular QTLs data have been widely used as an essential intermediate link of the non-coding disease risk variants to disease phenotypes. For example, more than four million common genetic variants (minor allele frequency >0.01)associated with the gene expression of more than 23,000 genes across 49 human tissues have been identified by Genotype-Tissue Expression (GTEx) Consortium (2), representing a valuable resource for the molecular interpretation of disease risk variants. eQTL catalogue (3) is another useful resource that includes uniformly processed expression QTLs (eQTLs) and splicing QTLs (sQTLs) based on, currently, 21 studies, and the data is still increasing. However, exploring and mining such a massive volume of xQTLs data remains computationally challenging. Firstly, xQTLs summaries typically contained millions of associations between genetic variants and molecular phenotypes, making retrieving all or a subset of xQTLs data of interest computationally expensive. Secondly, xQTLs data generated by different analysis pipelines with varying tools often did not correctly harmonize with rigorous quality control to boost statistical power and reduce false discovery in downstream analysis. Thirdly, currently, no tools can seamlessly annotate the xQTLs data of their underlying function.

Another challenge for analyzing xQTL data is that xQTLs are often integrated with GWAS summary statistics data for colocalization analysis, widely used to link potentially causal genes to GWAS risk loci. Several available tools can perform probabilistic colocalization analysis between xQTLs and GWAS summary statistics, including *Coloc* (4), *HyPrColoc* (5), *ColocQuiaL* (6), *ezQTL* (7), etc. Despite these tools being frequently used to identify disease-risk genes in many studies, many limitations remain. For instance, relying solely on a single colocalization tool may not identify reliable disease-related genes due to insufficient detection power, especially in cases where the sample size is small, or the causal variants have a relatively small effect size (8). In addition, all these tools only focus on statistical methods without considering the upstream data processing and downstream visualization of the results, making the colocalization analysis still computationally challenging. To our knowledge, no comprehensive analysis pipeline can seamlessly mine and analyze both xQTLs and GWAS data.

To address these challenges, we developed xQTLbiolinks, a user-friendly R package, as the first end-to-end bioinformatic tool for efficient mining and analyzing public and user-customized xQTLs data for the discovery of disease susceptibility genes. xQTLbiolinks offers the following unique and practical advantages: i) enables flexible accessing to xQTLs data from 23,839 samples across 75 human tissues or cell types and provides quality control and annotation modules for summary statistics data; ii) offers a robust colocalization pipeline that utilizes two popular colocalization methods to streamlining identity colocalized disease-associated genes. iii) compatible with external tools for downstream analysis and can flexibly generate detailed outputs and ready-to-publish figures. Our package is freely available at https://github.com/lilab-bioinfo/xQTLbiolinks.

## RESULTS

### xQTLbiolinks as a comprehensive tool for exploring and mining xQTLs data

xQTLbiolinks presents the first end-to-end solution for molecular QTL data mining and analyzing. Compared to previous tools, xQTLbiolinks provides comprehensive and versatile approaches to accessing and manipulating xQTLs summary data (**Table 1**). It streamlines the querying and retrieval of eQTL and sQTL data to meet user-customized demand from many xQTLs datasets in public repositories (i.e., GTEx and eQTL catalogue). Notably, it can also analyze user-customized xQTLs summary data. xQTLbiolinks is characterized by its exceptional adaptability and user-friendliness for the analysis of xQTLs data. Its key strengths can be attributed to the following factors: i) xQTLbiolinks was developed as a standardized R package which makes it easily accessible to users who are familiar with other R packages; ii) the utilization of xQTLs datasets in public repositories, i.e., largest atlas of human gene expression, eQTL and sQTL data from GTEx (2), uniformly processed xQTLs summary statistics across 75 distinct cell types and tissues from the eQTL Catalogue (3). xQTLbiolinks allows users to conveniently retrieve xQTLs data and metainformation for further analysis through gene names/IDs, tissue names, or genomic regions of interest. It also facilitates easy query and retrieval of either eQTLs or sQTLs for a specific gene or set of genes in particular genomic regions or all xQTLs for all genes in each tissue. Such flexibility saves running time and decreases the requirement of computational resources, thus taking full advantage of public and user-customized xQTLs data for GWAS integration of varying scales from candidate genes to genome-wide.

**Table 1.**
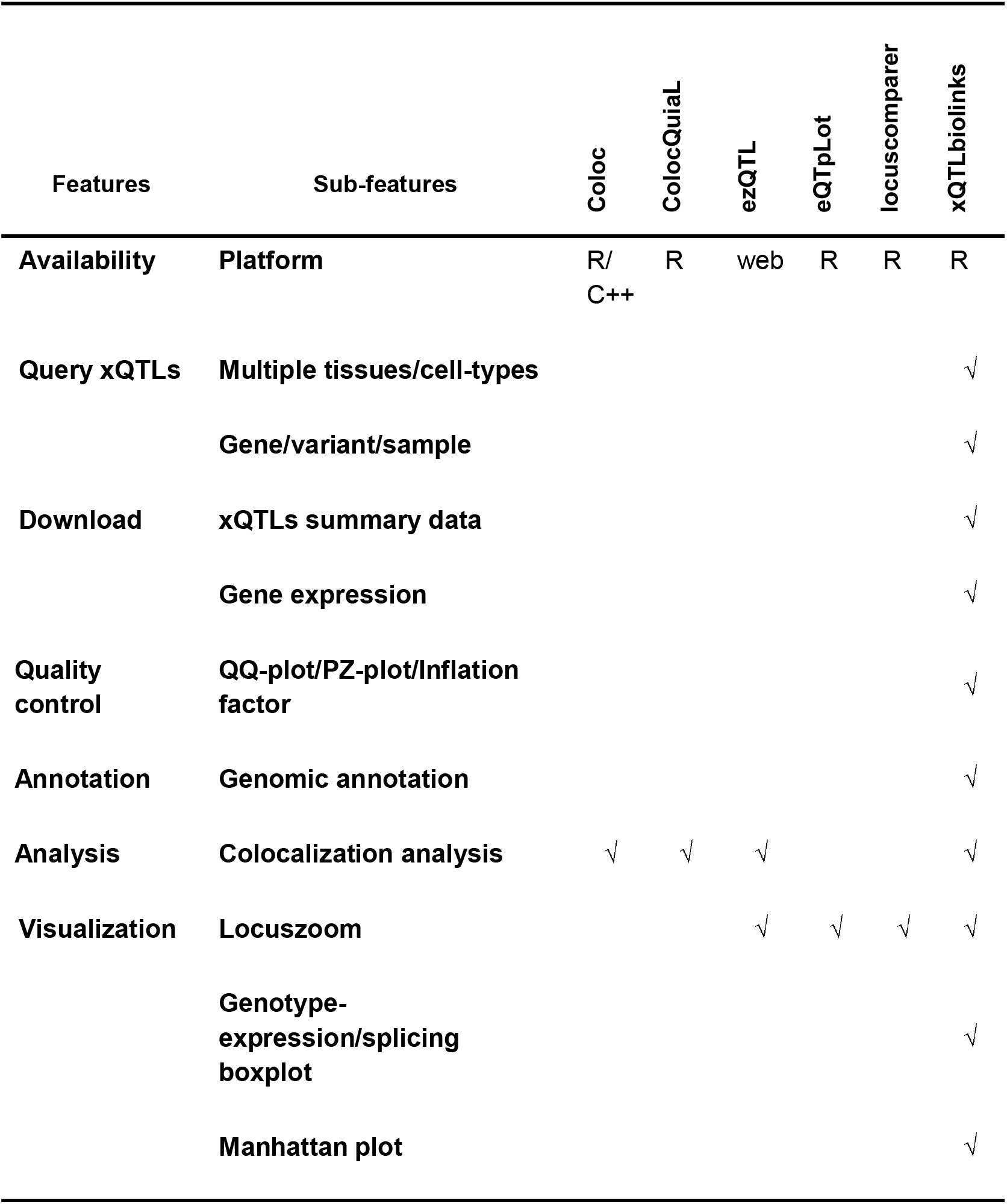
A comparison of different tools for retrieving and analysis of xQTLs data. The first two columns of the table represent features and detailed features for each tool, respectively. The cell checked with “√”indicates features that exist in the tool.

### xQTLbiolinks provides a robust colocalization module through integration with GWAS data

Colocalization is a powerful approach for integrating xQTLs and GWAS signals and has been widely used to identify novel disease susceptibility genes (9). This approach evaluates whether xQTLs and GWAS signals statistically share putative causal variants and can provide valuable insights into the genetic mechanisms underlying complex diseases. Besides, employing multiple colocalization methods can significantly improve the reliability of colocalized genes. Popular colocalization software such as Coloc and HyPrColoc are exclusively designed for only performing colocalization tests in a single condition(4,5), and visualization tools such as eQTpLot, and LocusCompare are solely used for visualization (10,11). xQTLbiolinks provides a comprehensive pipeline that can perform colocalization analysis across multiple tissues or cell types and handle upstream data processing and downstream visualization (**Fig. S1**). This framework offers a one-step solution of functions that can be used for quality control of GWAS significant variants, extraction of sentinel SNP, identification of trait genes, preparation of curated or user-customized xQTLs datasets, colocalization analysis, and visualizing the GWAS/xQTLs signals using locus zoom plot. Furthermore, to investigate the regulatory effect of colocalized xQTLs across multiple tissues/cell types, we have made a new plot by correlating xQTLs p-values with linkage disequilibrium (LD) bins.

### Case study 1: Quality control and functional characterization of breast cancer risk SNPs using xQTLbiolinks

We first downloaded the Breast cancer GWAS summary statistics from the literature, representing the largest GWAS study on breast cancer conducted on more than 80,000 individuals (12). We then performed a quality control analysis of the data. We first use *xQTLanno_calLambda* to estimate the p-values inflations, and it returns a lambda value of 1.147, indicating no strong population stratification. Then we evaluate the quality of GWAS data by examining whether the observed distribution of p-values follows the expected distribution under the null hypothesis of no association between the genetic variants and the disease using *xQTLvisual_qqplot*; we found a significant deviation from the diagonal line, which indicates potential variations from the null hypothesis that may result from true associations or LD (**Fig. 2a**). *xQTLvisual_PZplot* is further used to investigate the normality of the distribution of Z-scores derived from Beta and SE, which outputs the strong concordance between the observed p-values and those calculated from Z-scores (**Fig. 2b**). We further annotated all significant GWAS variants by genomic locations using *xQTLanno_genomic* (**Fig. 2c**). We also retrieved eQTLs and sQTLs summary data from GTEx Breast - Mammary tissue using *xQTLdownload_eqtlAllAsso* and *xQTLdownload_sqtlAllAsso functions*, respectively. In addition, we included our recently developed 3’UTR alternative polyadenylation quantitative trait loci (3’aQTLs) as a user-custom xQTLs dataset using *xQTLdownload_xqtlAllAsso*. We later performed similar quality control analyses for these xQTLs data and found no inflation or quality issues on these datasets. The genomic control inflation values of the xQTLs summary data are 1.149 (eQTLs), 1.049 (sQTLs), and 1.022 (3’aQTLs), and the corresponding QQ-plots are shown in **Fig. 2d-f**. We used *xQTLvisual_manhattan* to generate a Manhattan plot, which exhibits strong signals of associations across all chromosomes at a genome-wide level (**Fig. 2g**). By integrating with eQTL signals, we detected some significant loci that showed significant associations with disease susceptibility but also exerted regulatory influence on gene expression. For instance, rs8018155 was a breast cancer risk variant and an eQTL, as illustrated in **Fig. 2h**. This suggests that risk variant rs8018155 plays a crucial role in modulating the expression of gene *CCDC88C* and may have potentially important implications for breast cancer.

**Fig. 1.**
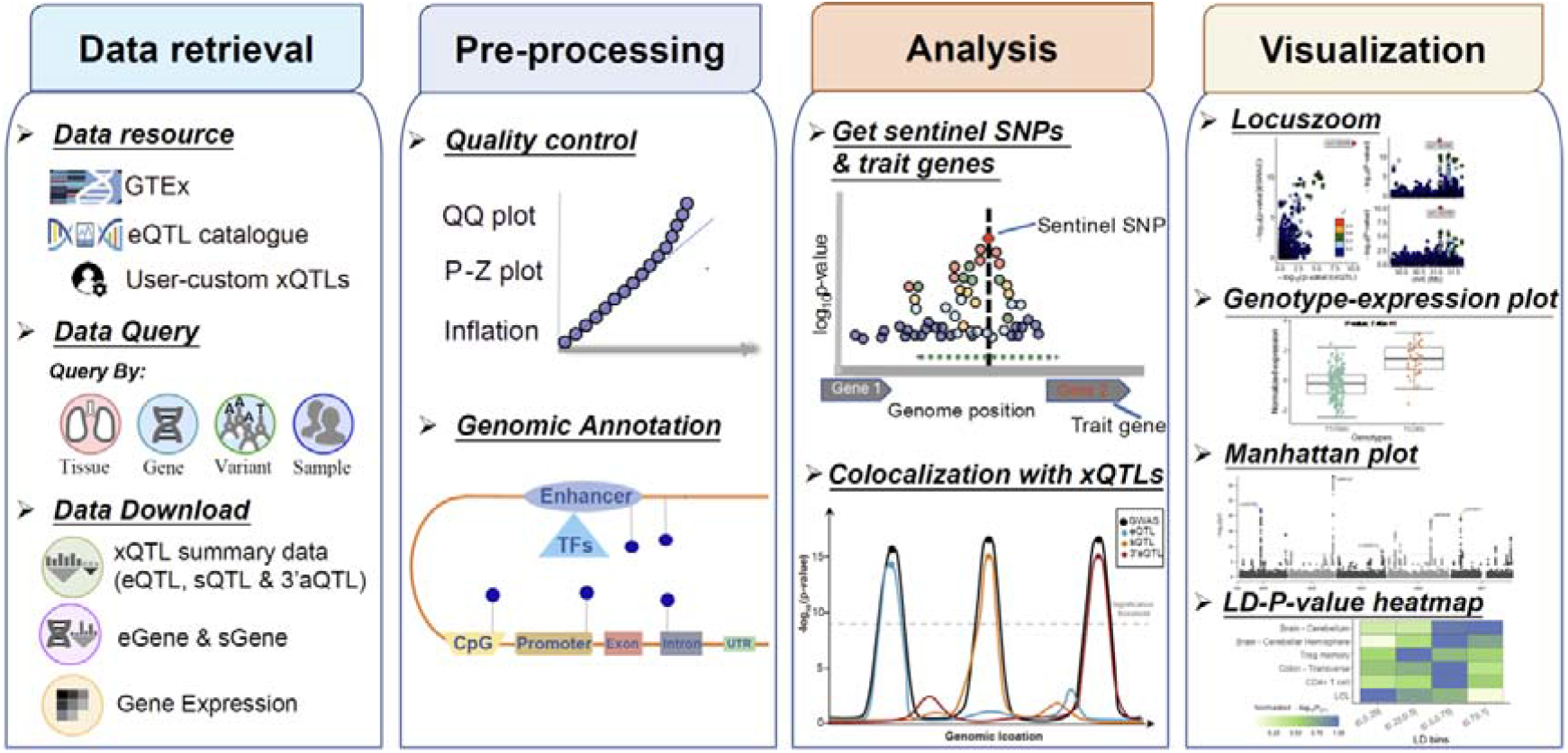
Overview of xQTLbiolinks data and functions, including four main function categories: Data retrieval, Pre-processing, Analysis, and Visualization.

**Fig. 2.**
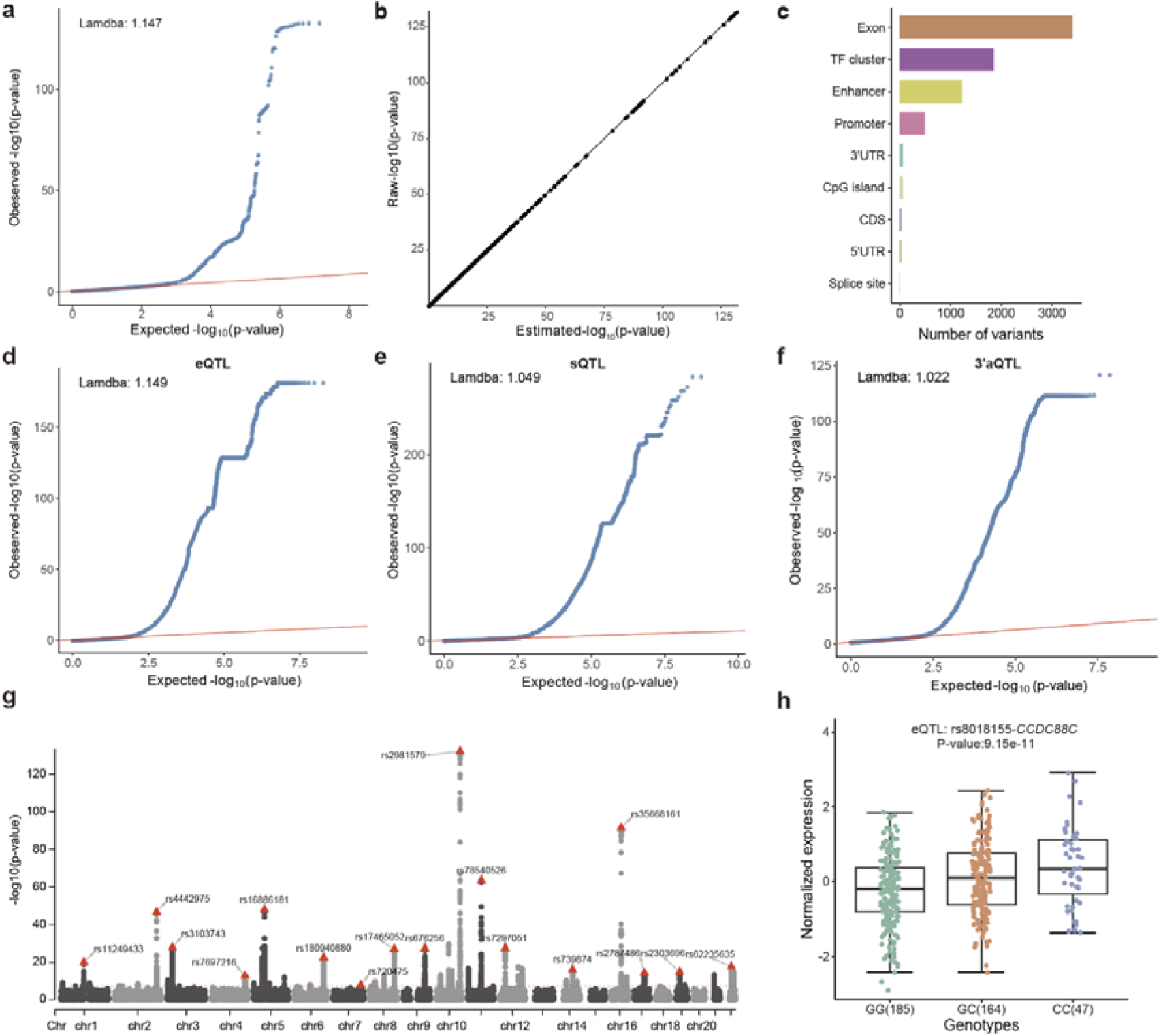
Quality control and annotation of breast cancer and xQTLs summary statistics data. (a). QQ-plot labeled with inflation factor. (b). PZ-plot, the x-axis stands for the normalized p-values estimated by the z-score derived from beta and standard error, and the y-axis stands for the raw normalized p-values. (c). Genomic annotation of significant breast cancer risk SNPs. (d) QQ-plot for eQTLs. (e) QQ-plot for sQTLs. (f) QQ-lot for 3’aQTLs. (g) Manhattan plot of the GWAS study of breast cancer. The strongest signals on each chromosome are labeled. (h) Boxplot of normalized expression of eQTL rs8018155–CCDC88C in the Breast - Mammary Tissue.

### Case study 2: Identification of prostate cancer susceptibility genes using xQTLbiolinks

Prostate cancer (PCa) is one of the most common cancers, the pathogenesis of which involves both heritable and environmental factors (13). The molecular events involved in the development or progression of PCa are still unclear. Here, we applied xQTLbiolinks to integrate the PCa GWAS dataset (14) with xQTLs data and aimed to identify putative causal variants and susceptibility genes associated with PCa. We first extracted 94 sentinel SNPs with *P*<5e-8 using *xQTLanalyze_getSentinelSnp* (**Table S1**). We then identified 835 genes for eQTLs, 1,676 genes for sQTLs, and 209 genes for 3’aQTLs using *xQTLanalyze_getTraits* **(Table S2)**. Later, for each trait gene, we analyzed the colocalization pattern between PCa risk variants and xQTLs using *xQTLanalyze_coloc* for eQTLs and *xQTLanalyze_coloc_diy* for sQTLs and 3’aQTLs. By default, two colocalization methods (Coloc and HyPrColoc) are used. The colocalization analysis returns four posterior probabilities corresponding to four different null hypotheses; notably, the posterior probability under hypothesis 4 (PPH4), representing the potential same causal variants shared by GWAS variants with xQTLs data, was used to define significant colocalization. Using a PPH4 threshold of 0.75, we identified 47 colocalized genes, including 27 eQTL colocalized genes, 17 sQTL colocalized genes, and seven 3’aQTL colocalized genes that are colocalized with 38 PCa risk loci. Among these co-localized genes, many have been previously reported to be associated with PCa susceptibility (**Table S3**). For example, the gene *MMP7*, which is strongly colocalized by eQTLs, encodes a member of the peptidase M10 family of matrix metalloproteinases and is involved in the breakdown of extracellular matrix in normal physiological processes. It has been evidenced that PCa can be promoted via MMP7-induced epithelial-to-mesenchymal transition by Interleukin-17 (15). and serum MMP7 levels could guide metastatic therapy for PCa (16). Notably, xQTLs colocalized genes are largely non-overlapped (**Fig. 3a**), such as *MMP7* was only significantly colocalized by eQTLs, *AGAP10P* was only colocalized by sQTLs rather than eQTLs and 3’aQTLs, suggesting the substantial added value of integrating multiple types of molecular QTLs data for disease susceptibility gene identification.

**Fig. 3.**
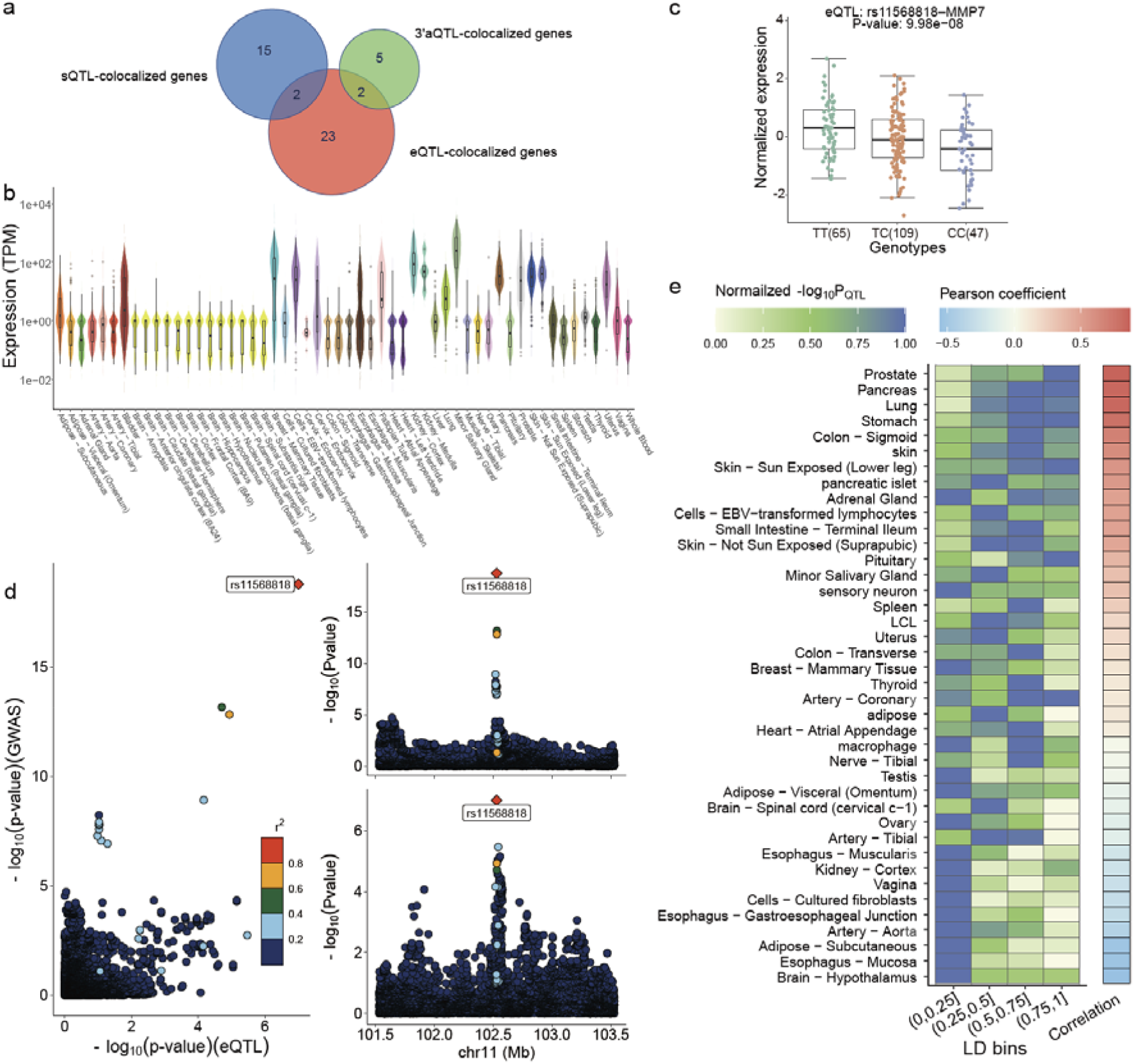
Integrative analysis of GWAS study of prostate cancer. (a) Venn plot of 47 xQTLs colocalized genes. (b) Gene expression levels of *MMP7* among 54 GTEx tissues. (c) Boxplot of normalized expression of eQTL rs11568818–MMP7 in the prostate. (d) Distribution of GWAS and eQTL signals within a genome region of *MMP7*. (e) heatmap of eQTL (rs11568818–MMP7) p-values in different LD bins across 40 tissues/cell types. The y-axis represents the tissues/cell types, and the x-axis represents LD bins. The green-blue color bar indicates the median normalized eQTL p-values in different LD bins. The blue-red color bar represents the Pearson correlation between normalized eQTL p-values and values of r-squared of LD.

To understand the colocalized results, we first visualize the distribution of *MMP7* gene expression across 49 GTEx tissues by function *xQTLvisual_geneExpTissues*. As shown in **Fig. 3b**, *MMP7* has a relatively high expression level in the prostate and relevant tissues, indicating a potential essential role in these tissues. The distribution of *MMP7* expression level in prostate tissue stratified by the genotype of the lead SNP was also presented by function *xQTLvisual_eqtlExp* (**Fig. 3c**). Then, we visualized the *MMP7* colocalized signals by functions *xQTLvisual_locusZoom*, which reveals a high correlation between GWAS variants and *MMP7* eQTLs (**Fig. 3d**). To further compare with other molecular QTLs, *xQTLvisual_locusZoom* have also implemented the functions to integrate and plot multiple xQTLs data (**Fig. 2a**).

xQTLbiolinks is highly compatible, allowing for seamless integration of its outputs with external packages. For example, we performed gene ontology (GO) enrichment analyses on the eQTL colocalized genes with external package clusterProfiler (11). Cancer-related GO terms are significantly enriched, including “positive regulation of T cell receptor signaling pathway”, “Execution phase of apoptosis, “and “DNA replication checkpoint signaling”. Moreover, we can also perform co-expression analysis using the corrplot package (**Fig. 2b**) (17). To investigate the regulatory sharing of colocalized variants across multiple tissues, we implemented *xQTLvisual_coloc* to visualize the correlation between p-values of xQTLs LD across numerous tissues/cell types (**Fig. 3e**). We observed that prostate tissues showed the strongest correlation indicating the heatmap can reveal the potential disease-relevant tissues.

## DISCUSSION

Molecular Quantitative Trait Loci is a crucial step towards better understanding the effects of non-coding genetic variants on genes, pathways, and their function mechanism and serve as an essential link between genotype and disease phenotype. Although many xQTLs summary statistics are available, mining and analyzing these xQTLs data remains several major bioinformatic challenges, such as data retrieval, quality control, and pre-processing, which are essentially required steps to promote the reproducible use of xQTLs resources and accelerate disease susceptibility genes identification remains challenging. Here, we developed xQTLbiolinks, which is motivated by TCGAbiolinks that provides several useful functions to search, download and prepare TCGA samples for data analysis (18). Here, xQTLbiolinks aims to “link” xQTLs data to disease genomics research by providing flexible interfaces to allow users to access xQTLs data from GTEx and eQTL catalogue without having to navigate through different data portal sites or download whole tables of millions of xQTLs associations. The current version of xQTLbiolinks provides access to over 4 million eQTLs and sQTLs in public servers. It enables manipulating user-customized xQTLs data, such as our recently developed 3’UTR alternative polyadenylation QTLs (3’aQTLs) (19). Our plan includes collecting more data resources, such as the eQTLGen and MetaBrain project (20,21).

In addition, colocalization analysis is a powerful approach that has been widely used to identify new susceptibility genes in disease analysis by integrating xQTLs and GWAS signals. However, current colocalization tools have limitations in that they only focus on colocalization methods without considering the whole analysis pipeline and the effects of multiple tissue or cell types. Employing several colocalization tools in the context of numerous tissue or cell types is a superior option to enhance the robustness and reliability of the findings (8). To facilitate the identification of robust susceptibility genes in colocalization analysis, xQTLbiolinks provides a comprehensive pipeline that employs two popular colocalization methods and handles upstream data processing and downstream visualization of results in a one-step solution.

xQTLbiolinks is a scalable tool that facilitates integrating and utilizing external tools or packages. We will also actively maintain xQTLbiolinks and respond to user inquiries. As the first end-to-end bioinformatic framework for mining and analyzing xQTL data for discovering disease susceptibility genes, it will make significant contributions toward our understanding of human non-coding variants, thus promoting a better understanding of disease etiologies.

## MATERIAL AND METHODS

### Implementation of xQTLbiolinks

xQTLbiolinks, a user-friendly R package under the General Public License (GPLv3) license, can be installed in any operating system supporting R through the general function install.packages(“xQTLbiolinks”) from the Comprehensive R Archive Network (CRAN: https://cran.r-project.org/). All functions of xQTLbiolinks are available by standard R commands to manipulate xQTLs and GWAS data after installation and loading of the package. A comprehensive user manual introducing all functions and corresponding usages can be found in the Github repository (https://github.com/lilab-bioinfo/xQTLbiolinks). Briefly, xQTLbiolinks implements four modules: data retrieval, pre-processing, colocalization analysis, and data visualization (**Fig. 1**). The functions and outputs in xQTLbiolinks are compatible with other functions and packages. Users can perform customized analysis with external R packages, including functional enrichment analysis using clusterProfiler (11), differential expression analysis using DESeq2 and edgeR (22,23), weighted Correlation Network Analysis using WGCNA (24), etc.

### Data retrieval

The data retrieval module enables users to query and download publicly available xQTLs summary data. The current version supports xQTLs data from 23,839 samples across 75 human tissues/cell types. Two commands, *xQTLquery* and *xQTLdownload*, provide flexible user interfaces to query and download xQTLs data by tissue, gene, SNP, or combination. For example, specifying a specific tissue name to the *xQTLquery* function will display all xQTLs data associated with the tissue; users can also focus only on xQTLs related to one single gene by specifying gene id to the function. *xQTLquery* is the primary function that executes the query of entities in xQTLs data, including genes, variants, and samples. At the same time, *xQTLdownload* allows users to retrieve xQTLs data with tailored demands, including eGenes/sGenes, associations between variants and expression (eQTLs) and splicing (sQTLs), normalized gene expressions, and linkage disequilibrium of xQTLs in specified genes. All retrieved xQTLs data can be handled by *xQTLanalyze, xQTLvisual*, and their sub-functions.

### Data pre-processing

xQTLbiolinks allows users to detect possible inflation or deflation of test statistics due to population stratification, genotyping errors, or other sources of bias for GWAS/QTL summary statistics datasets, *xQTLvisual_qqPlot* and *xQTLanno_calLambda* can plot quantile-quantile plot (QQ plot) and calculate genomic control inflation factor, respectively. xQTLvisual_PZPlot plots the concordance correlation of observed P-values and P-values calculated from the Z-score derived from Beta (representing effect size) and SE (representing standard error of effect size). Besides, *xQTLanno_genomic* enables users to functionally annotate all variants in GWAS/xQTLs datasets by genome location, including intron, exon, 3 ‘UTR, 5’UTR, promoter, CDS, splicing site, intergenic region, transcription factor binding sites (TFBS), CpG Island, and enhancer.

### Colocalization analysis

xQTLbiolinks implements a pipeline that contains three subfunctions to perform colocalization analysis following the steps (**Fig. S1**): (i) *xQTLanalyze_getSentinelSnp* extracts sentinel SNP, which represents the most prominent signal in a specific genome region, from GWAS summary statistics data. By default, it selects the variants with a p-value less than 5 × 10^−8^ within 1 million base pairs (1Mbp). (ii) *xQTLanalyze_getTraits* identify trait genes nearby sentinel SNPs within a 1Mb region by default. (iii) *xQTLanalyze_coloc* and xQTLanalyze_coloc_diy perform two commonly used colocalization methods, Coloc (4) and HyPrColoc (5), for each trait gene.

### Visualization

The visualization module contains a main function *xQTLvisual* and several sub-functions that allow users to visualize the results with publication-ready plots, such as heatmap, boxplot, scatter plot, and locus zoom plot. *xQTLvisual_genesExp* and *xQTLvisual_geneExpTissues* are used to plot the distribution of gene expression for the queried gene(s). Sub-functions *xQTLvisual_eqtlExp* and *xQTLvisual_sqtlExp* can plot the association between genotypes and molecular phenotypes with a grouped boxplot by genotypes. xQTLbiolinks also contains three sub-functions *xQTLvisual_locusZoom* and *xQTLvisual_locusCompare, xQTLvisual_coloc* visualizing the colocalization results. The first two subfunctions can plot publication-ready locus zoom on specific GWAS and xQTLs signals. And xQTLvisual_coloc can visualize the regulatory sharing effects of colocalized variants across multiple tissues or cell types.

## DATA AVAILABILITY

xQTLbiolinks and the standard manual (version 1.5.2) are publicly available at CRAN https://cran.r-project.org/web/packages/xQTLbiolinks. The source codes of the latest version of xQTLbiolinks can be found in Github repository https://github.com/lilab-bioinfo/xQTLbiolinks.

## AUTHOR CONTRIBUTIONS

Ruofan Ding: Conceptualization, Formal analysis, Methodology, Validation, Writing—original draft. Xudong Zou: Conceptualization, Formal analysis, Validation, Writing—review & editing. Xuelian Ma & Yangmei Qin: Formal analysis, Validation. Hui Chen: Writing-review. Chen Yu: Writing-review & editing. Gao Wang: Methodology, Writing—review & editing. Lei Li: Conceptualization, Methodology, Validation, Writing—review & editing.

## ACKNOWLEDGEMENTS

We acknowledge all members of the Li lab for constructive discussions and help. We also thank Qin Wang at Shenzhen Bay Laboratory supercomputing center for high-computing support.

## TABLE AND FIGURES LEGENDS

**Fig. S1.**
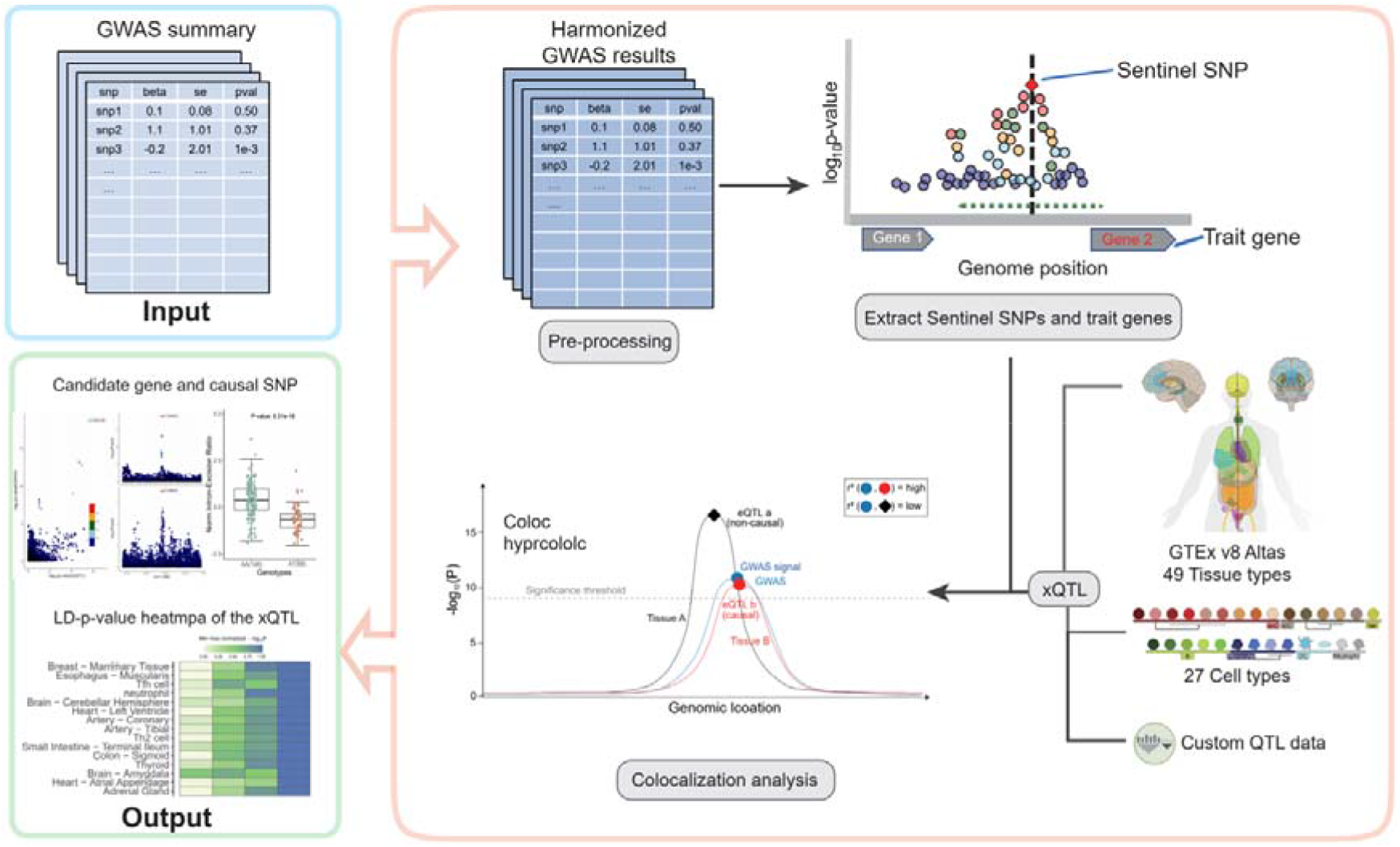
The processing flow of colocalization analysis in xQTLbiolinks. The framework takes in a GWAS summary statistics dataset as input. The output is a list of colocalized genes and causal SNPs, which can be visualized through locus zoom, boxplot, and heatmap. Data pre-processing and analysis are conducted in the following steps: quality control, removal of missing values and duplicates, extracting sentinel SNPs and trait genes, retrieval of xQTLs summary statistics dataset, and colocalization analysis.

**Fig. S2.**
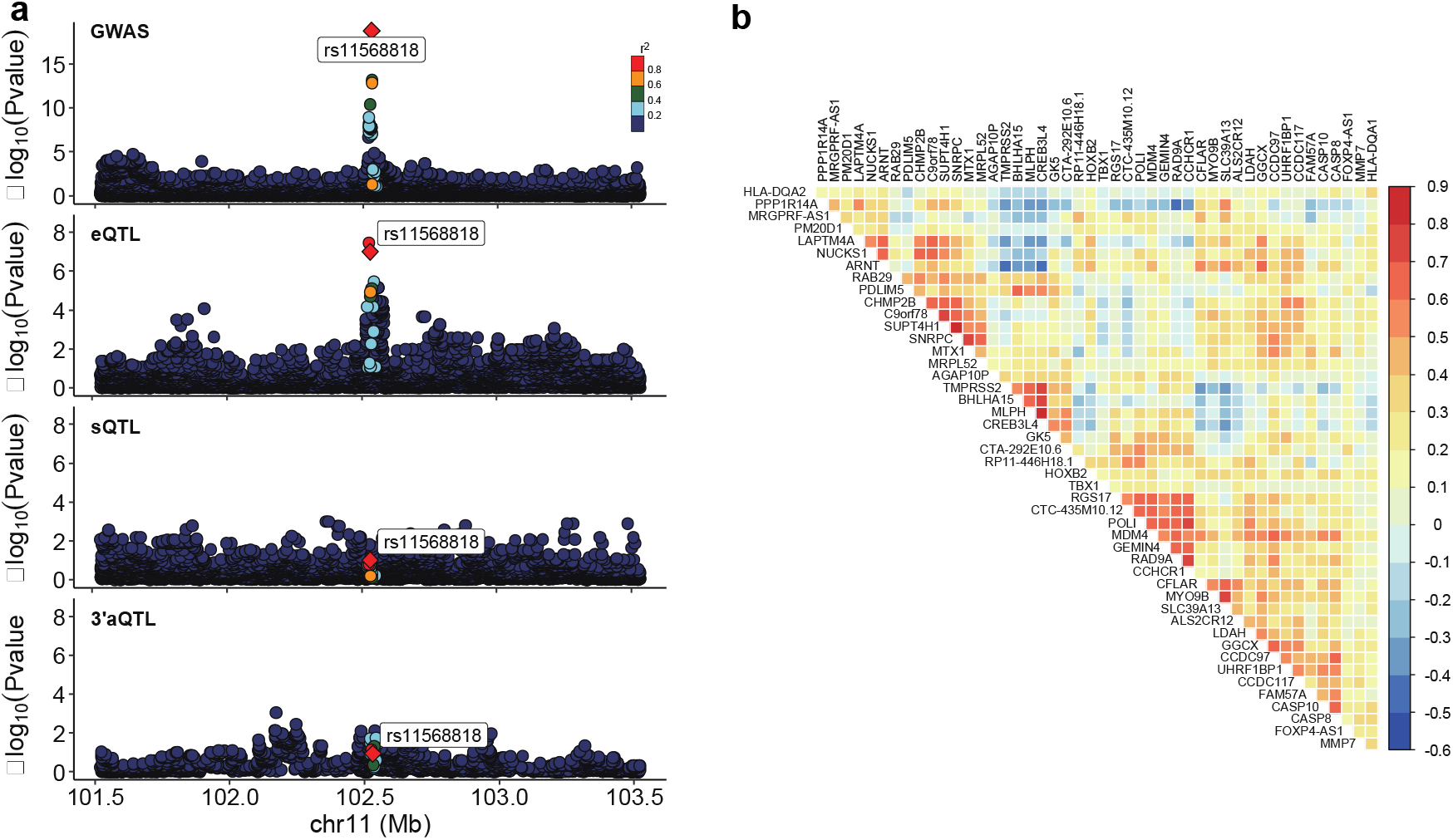
(a). Colocalization of signals of GWAS and xQTLs (eQTLs, sQTLs, and 3’aQTLs) for gene *MMP7* with prostate cancer-associated loci. (b). Co-expression analysis of 47 xQTL colocalized genes.

## Reference

1. Hormozdiari, F., Gazal, S., van de Geijn, B., Finucane, H.K., Ju, C.J., Loh, P.R., Schoech, A., Reshef, Y., Liu, X., O’Connor, L. et al. (2018) Leveraging molecular quantitative trait loci to understand the genetic architecture of diseases and complex traits. Nat Genet, 50, 1041–1047.

2. Consortium, G.T. (2020) The GTEx Consortium atlas of genetic regulatory effects across human tissues. Science, 369, 1318–1330.

3. Kerimov, N., Hayhurst, J.D., Peikova, K., Manning, J.R., Walter, P., Kolberg, L., Samovica, M., Sakthivel, M.P., Kuzmin, I., Trevanion, S.J. et al. (2021) A compendium of uniformly processed human gene expression and splicing quantitative trait loci. Nat Genet, 53, 1290–1299.

4. Giambartolomei, C., Vukcevic, D., Schadt, E.E., Franke, L., Hingorani, A.D., Wallace, C. and Plagnol, V. (2014) Bayesian test for colocalisation between pairs of genetic association studies using summary statistics. PLoS Genet, 10, e1004383.

5. Foley, C.N., Staley, J.R., Breen, P.G., Sun, B.B., Kirk, P.D.W., Burgess, S. and Howson, J.M.M. (2021) A fast and efficient colocalization algorithm for identifying shared genetic risk factors across multiple traits. Nat Commun, 12, 764.

6. Chen, B.Y., Bone, W.P., Lorenz, K., Levin, M., Ritchie, M.D. and Voight, B.F. (2022) ColocQuiaL: a QTL-GWAS colocalization pipeline. Bioinformatics, 38, 4409–4411.

7. Zhang, T., Klein, A., Sang, J., Choi, J. and Brown, K.M. (2022) ezQTL: A Web Platform for Interactive Visualization and Colocalization of QTLs and GWAS Loci. Genomics Proteomics Bioinformatics, 20, 541–548.

8. Hukku, A., Pividori, M., Luca, F., Pique-Regi, R., Im, H.K. and Wen, X. (2021) Probabilistic colocalization of genetic variants from complex and molecular traits: promise and limitations. Am J Hum Genet, 108, 25–35.

9. Hormozdiari, F., van de Bunt, M., Segre, A.V., Li, X., Joo, J.W.J., Bilow, M., Sul, J.H., Sankararaman, S., Pasaniuc, B. and Eskin, E. (2016) Colocalization of GWAS and eQTL Signals Detects Target Genes. Am J Hum Genet, 99, 1245–1260.

10. Drivas, T.G., Lucas, A. and Ritchie, M.D. (2021) eQTpLot: a user-friendly R package for the visualization of colocalization between eQTL and GWAS signals. BioData Min, 14, 32.

11. Liu, B., Gloudemans, M.J., Rao, A.S., Ingelsson, E. and Montgomery, S.B. (2019) Abundant associations with gene expression complicate GWAS follow-up. Nat Genet, 51, 768–769.

12. Michailidou, K., Beesley, J., Lindstrom, S., Canisius, S., Dennis, J., Lush, M.J., Maranian, M.J., Bolla, M.K., Wang, Q., Shah, M. et al. (2015) Genome-wide association analysis of more than 120,000 individuals identifies 15 new susceptibility loci for breast cancer. Nat Genet, 47, 373–380.

13. Braga-Basaria, M., Dobs, A.S., Muller, D.C., Carducci, M.A., John, M., Egan, J. and Basaria, S. (2006) Metabolic syndrome in men with prostate cancer undergoing long-term androgen-deprivation therapy. J Clin Oncol, 24, 3979–3983.

14. Schumacher, F.R., Al Olama, A.A., Berndt, S.I., Benlloch, S., Ahmed, M., Saunders, E.J., Dadaev, T., Leongamornlert, D., Anokian, E., Cieza-Borrella, C. et al. (2018) Association analyses of more than 140,000 men identify 63 new prostate cancer susceptibility loci. Nat Genet, 50, 928–936.

15. Zhang, Q., Liu, S., Parajuli, K.R., Zhang, W., Zhang, K., Mo, Z., Liu, J., Chen, Z., Yang, S., Wang, A.R. et al. (2017) Interleukin-17 promotes prostate cancer via MMP7-induced epithelial-to-mesenchymal transition. Oncogene, 36, 687–699.

16. Tregunna, R. (2020) Serum MMP7 levels could guide metastatic therapy for prostate cancer. Nat Rev Urol, 17, 658.

17. Taiyun Wei, V.S. (2021) R package ‘corrplot’: Visualization of a Correlation Matrix.

18. Colaprico, A., Silva, T.C., Olsen, C., Garofano, L., Cava, C., Garolini, D., Sabedot, T.S., Malta, T.M., Pagnotta, S.M., Castiglioni, I. et al. (2016) TCGAbiolinks: an R/Bioconductor package for integrative analysis of TCGA data. Nucleic Acids Res, 44, e71.

19. Li, L., Huang, K.L., Gao, Y., Cui, Y., Wang, G., Elrod, N.D., Li, Y., Chen, Y.E., Ji, P., Peng, F. et al. (2021) An atlas of alternative polyadenylation quantitative trait loci contributing to complex trait and disease heritability. Nat Genet, 53, 994–1005.

20. de Klein, N., Tsai, E.A., Vochteloo, M., Baird, D., Huang, Y., Chen, C.Y., van Dam, S., Oelen, R., Deelen, P., Bakker, O.B. et al. (2023) Brain expression quantitative trait locus and network analyses reveal downstream effects and putative drivers for brain-related diseases. Nat Genet, 55, 377–388.

21. Vosa, U., Claringbould, A., Westra, H.J., Bonder, M.J., Deelen, P., Zeng, B., Kirsten, H., Saha, A., Kreuzhuber, R., Yazar, S. et al. (2021) Large-scale cis- and trans-eQTL analyses identify thousands of genetic loci and polygenic scores that regulate blood gene expression. Nat Genet, 53, 1300–1310.

22. Love, M.I., Huber, W. and Anders, S. (2014) Moderated estimation of fold change and dispersion for RNA-seq data with DESeq2. Genome Biol, 15, 550.

23. Robinson, M.D., McCarthy, D.J. and Smyth, G.K. (2010) edgeR: a Bioconductor package for differential expression analysis of digital gene expression data. Bioinformatics, 26, 139–140.

24. Langfelder, P. and Horvath, S. (2008) WGCNA: an R package for weighted correlation network analysis. BMC Bioinformatics, 9, 559.

